# Mucosal Inflammation Shapes Human Neutrophil States in Tissue and Circulation

**DOI:** 10.64898/2026.03.21.713286

**Authors:** David Fraser, Vasileios I. Theofilou, Teresa Greenwell-Wild, Laurie Brenchley, Eleni Kanasi, Chen Wang, Niki M. Moutsopoulos

## Abstract

The oral mucosa is a prototypical human barrier reliant on neutrophils for homeostasis, as both neutrophil deficiency and excessive activation are linked to immunopathology. Yet, whether neutrophils acquire tissue-specific states in health or disease remains unclear. We incorporated single-cell RNA sequencing, spectral flow cytometry, and spatial proteomics across tooth-associated oral mucosa (gingiva) and interconnected compartments of blood and oral cavity to define neutrophil tissue specification in healthy individuals and patients with periodontitis, a neutrophil-dominated inflammatory disease. In health, mucosal neutrophils adopt discrete immunoregulatory states despite constant microbial exposure and mechanical injury. Periodontitis disrupts these programs through infiltration of blood-like neutrophil subsets, increased transcriptional noise, and heightened effector activation. Strikingly, oral inflammation systemically imprints on circulating neutrophils, marked by the expansion of a Rho-GTPase regulatory program that is shared across diverse human inflammatory conditions. Together, these findings establish a framework for understanding how localized tissue inflammation affects both neutrophil plasticity at barrier surfaces and conditioning of systemic neutrophil states with broad implications for inflammatory disease pathogenesis.

## Introduction

Neutrophils are the most abundant immune cell and are well-recognized as pillars of the innate immune system(1, 2). Billions of neutrophils exit the bone marrow each day, poised to migrate to sites of infection and injury and perform critical functions for antimicrobial defense and wound clearance(3–5). Beyond these well-recognized roles, work over the past decades has illuminated diverse neutrophil functions ranging from mediating immunopathology and triggering autoimmunity to facilitating immunomodulation, inflammatory resolution, and tissue repair(6–8). Key in the recognition of these pleotropic roles has been seminal work, performed mostly in experimental model systems, leveraging advances in microscopy and single cell sequencing to identify molecular programs driving neutrophil development, aging and tissue adaptation programs which, in turn, can dictate divergent neutrophil responses in specific contexts(9–13). Despite this expanded understanding of neutrophil states and functionality in various models, representation of neutrophils in human mucosal atlases remains sparse(14–17), leaving the tissue specification and functionality of human neutrophils far less understood.

A question of great importance to human health is the function of neutrophils within human barrier tissues. Barriers of the mammalian host are constantly exposed to the outside environment and are primary sites for infection and injury, necessitating constant immune patrolling and inflammatory regulation. The oral mucosa is a prototypic barrier tissue where neutrophil functionality is critical for homeostasis(18). Here, neutrophils constantly transmigrate through blood vessels into the tooth-associated mucosa (gingiva), even in the setting of pristine clinical health, and pass into the mouth via a particularly penetrable epithelial barrier at the interface of the gingiva, the tooth, and the tooth-associated bacterial biofilms(19–21). In fact, one of the earliest mentions of phagocytotic cells in the scientific literature was a description of neutrophils isolated from the oral cavity (“salivary corpuscles”) in 1869(22, 23). In the more recent past, neutrophil counts in the oral cavity were employed for the diagnosis of patients with neutrophil recruitment defects, prior to the advent of genetic testing(24). The critical role of neutrophils in the oral mucosa becomes most evident through the study of patients with single gene disruptions linked to neutrophil development and trafficking who present with dominant oral clinical phenotypes, including severe, early-onset periodontitis and recurrent oral ulcerations(25–27).

While neutrophils are critical for oral mucosal homeostasis, excessive neutrophil recruitment and activation has also been linked to immunopathology in common (non-monogenic) forms of periodontitis(28–31), a microbiome-triggered disease of the gingiva defined by chronic mucosal inflammation and resorption of the underlying bone(32, 33). Periodontitis has been linked to systemic inflammatory diseases such as diabetes, rheumatoid arthritis, and cardiovascular disease, among others(34, 35), underscoring the greater consequences of the disease beyond its oral manifestations. Whether homeostatic versus inflammatory roles at the oral mucosa are related to the sheer balance of neutrophil numbers or are a consequence of distinct neutrophil subsets mediating specific functions in health or disease is a key question for the field of mucosal immunity.

To understand human neutrophil specification within the mucosal environment, we conducted a meticulously designed human study, acquiring neutrophils across tissue compartments (blood, gingiva, and the oral cavity) from healthy volunteers and patients with severe periodontitis. We employed a muti-omics approach, combining single-cell RNA sequencing (scRNA-seq) with spectral flow cytometry and spatial proteomics to define neutrophil subsets and states with spatial context in oral mucosal health and disease. Our work provides a resource towards defining human neutrophil subsets and their adaptation in the mucosal tissue microenvironment and reveals imprinting of mucosal inflammation on local tissue and systemic neutrophils in the context of the oral disease, periodontitis.

## Results

### Neutrophils and Neutrophil Extracellular Traps are Integral Components of the Tooth-Associated Mucosal Barrier

In healthy individuals, neutrophils continuously migrate into the gingiva and across the permeable tooth-associated epithelium (TAE) into the oral cavity, with infiltration increasing significantly in periodontitis(15, 36) (**Figure 1A**). This constitutive neutrophil presence at the TAE and the accessibility of the oral cavity renders the gingiva an optimal site for investigating human neutrophil specification and function within homeostatic and inflamed mucosal environments.

**Figure 1.**
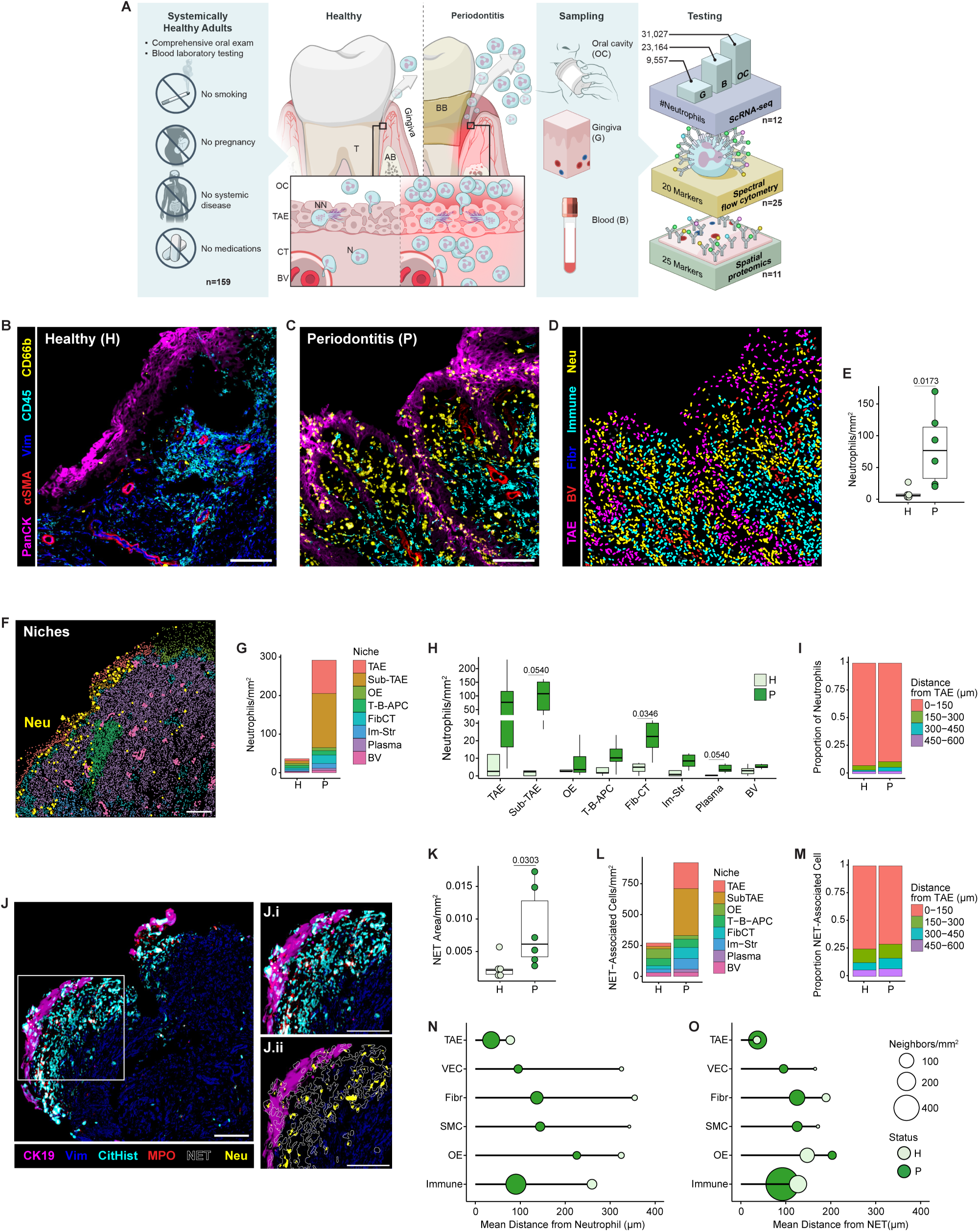
Neutrophils and Neutrophil Extracellular Traps are Integral Components of the Tooth-Associated Mucosal Barrier. A) Left: Clinical study design, inclusion/exclusion criteria. Center: schema of the gingiva and surrounding tissues. T: Tooth, AB: alveolar bone, BB: bacterial biofilm, N: neutrophil, NN: neutrophil NETs, BV: blood vessels, CT: connective tissue, TAE: tooth-associated epithelium, OC: oral cavity (OC). Right: sampling sites and testing modalities. B-C) Representative images of healthy (H) and periodontitis (P) gingiva stained with pan cytokeratin (PanCK, epithelium), α-smooth muscle actin (αSMA, BV), vimentin (Vim, stromal cells), CD45 (immune cells), and CD66b (neutrophils, Neu). D) Nuclear segmentation-based annotation of (D). E) Neutrophil numbers within H and P gingiva normalized by tissue area. F) Representative image of gingival niches with neutrophils marked in yellow. G) Distribution of neutrophils across niches in H and P. OE: oral epithelium, FibCT: fibrous connective tissue, Im-Str: immune-stromal. H) Neutrophils per niche normalized by tissue area. I) Proportion of neutrophils within the given distance from the TAE in H or P gingiva. J) Representative image of gingiva showing neutrophil extracellular traps (NETs) identified as citrullinated histone H3 (CitHist) and myeloperoxidase (MPO) structures with insets showing staining adjacent to the CK19 positive TAE and associated neutrophils (CD66+). K) NET area normalized by tissue area. L) Distribution of NET-associated cells by tissue niche. M) Proportion NET-associated cells within the given distance from the TAE. N and O) Number of neighbors and distances between neutrophil (N) or NETs (O) and epithelial cells (TAE or OE), stromal cells (VEC: vascular endothelial cells, Fibr: fibroblasts, SMC: smooth muscle cells), or immune cells. Circles represent the mean number of neighbors, defined as cells within 18 μm (30 pixels) of a neutrophil or NET and normalized by tissue area, and lines indicates the mean distance between neutrophils or NETs and the given cell type (n=11). Scale bars are 100 μm. Symbols represent individual subjects in E and K and mean values in N and O. Wilcoxon-signed rank test was used for E, H, and K with Benjamini-Hochburg adjustment for multiple comparisons applied in H. Lines with numbers represent p-values in E and K and adjusted p-values in H.

To investigate neutrophils at this barrier, we prospectively recruited healthy individuals and patients with severe, untreated periodontitis, excluding those with known comorbidities, and collected paired samples from blood and the oral cavity, along with gingival biopsies or surgical specimens. Using a multi-omics approach, including scRNA-seq, spectral flow cytometry, and spatial proteomics, we sought to characterize neutrophil subpopulations and states across tissue compartments in health and disease. Our dataset included 63,748 neutrophil transcriptomes from 12 subjects, surface marker analysis of 3,541,423 neutrophils from 25 subjects, and spatial analysis of 12,532 gingival neutrophils (**Figure 1A, Table S1**).

Spatial proteomics revealed neutrophils at low densities in healthy gingiva and significantly elevated numbers in periodontitis, aligning with previous findings(21, 37) (**Figure 1B-E, Figure S1A-B**). Cellular neighborhood (niche)-based analysis confirmed the presence of neutrophils within the TAE in health and disease (**Figure 1B-C, Figure S1C-D**). Periodontitis was associated with increased neutrophil infiltration in the TAE and the expansion of a neutrophil-rich sub-TAE niche (**Figure 1F-H, Figure S1C-E**). Smaller but significant increases in neutrophil numbers were also detected in deeper connective tissue niches (**Figure 1H**). Overall, the spatial distribution of neutrophils was strongly driven by proximity to the TAE, with the majority of neutrophils located within 150 µm of this structure in both healthy and diseased tissue (**Figure 1I**).

As previous work has associated neutrophil extracellular traps (NETs) with periodontitis(31, 38), we evaluated mucosal tissues for the presence of citrullinated histone (CitHist)+/myeloperoxidase (MPO)+ structures. NETs were observed in both healthy and diseased gingiva (**Figure 1J**) and NET-covered area was significantly increased in periodontitis (**Figure 1K**). NETs localized primarily within TAE and sub-TAE niches with a reduced presence in deeper regions, mirroring the distribution of neutrophils (**Figure 1lL-M, Figure S1F**).

To assess potential neutrophil interactions within the tissue microenvironment, we examined neutrophil and NET proximity to other cell types. Both were closely associated with TAE cells regardless of disease status, consistent with their localization at this specific epithelium (**Figure 1N-O**). Beyond the TAE, neutrophils and NETs showed the strongest proximity with immune cells, particularly in periodontitis (**Figure 1N-O, Figure S1E**), suggesting the potential for neutrophil- and NET-immune cell interactions in this environment.

Together, these findings confirm that neutrophils are a consistent feature of the gingiva in both health and disease and identify a spatially distinct pattern of neutrophil infiltration during chronic inflammation. The presence of neutrophils proximal to the TAE suggests they may help secure the permeable border of this barrier tissue, in part, through NET structures. Further, the close association between neutrophils, NETs, and immune cells suggests a role for neutrophils in modulating immune responses at this critical interface.

### Cell surface markers reflect activation of neutrophil effector functions in the oral environment

We next sought to characterize neutrophils across tissue compartments via spectral flow cytometry to identify how migration into gingiva and exit into the oral cavity affects neutrophil states (**Figure S2A**). In the oral cavity, neutrophils were reliably detected in healthy subjects and increased in number by ∼8-fold in periodontitis (**Figure 2A**). In circulation, we detected few low-density neutrophils (LDNs) in either healthy or periodontitis subjects’ blood (**Figure 2B**) and minimal immature neutrophils (CD10–) were identified in any compartment (**Figure 2C**), indicating that oral neutrophils are primarily derived from mature normal-density neutrophils in both health and chronic oral inflammation.

**Figure 2.**
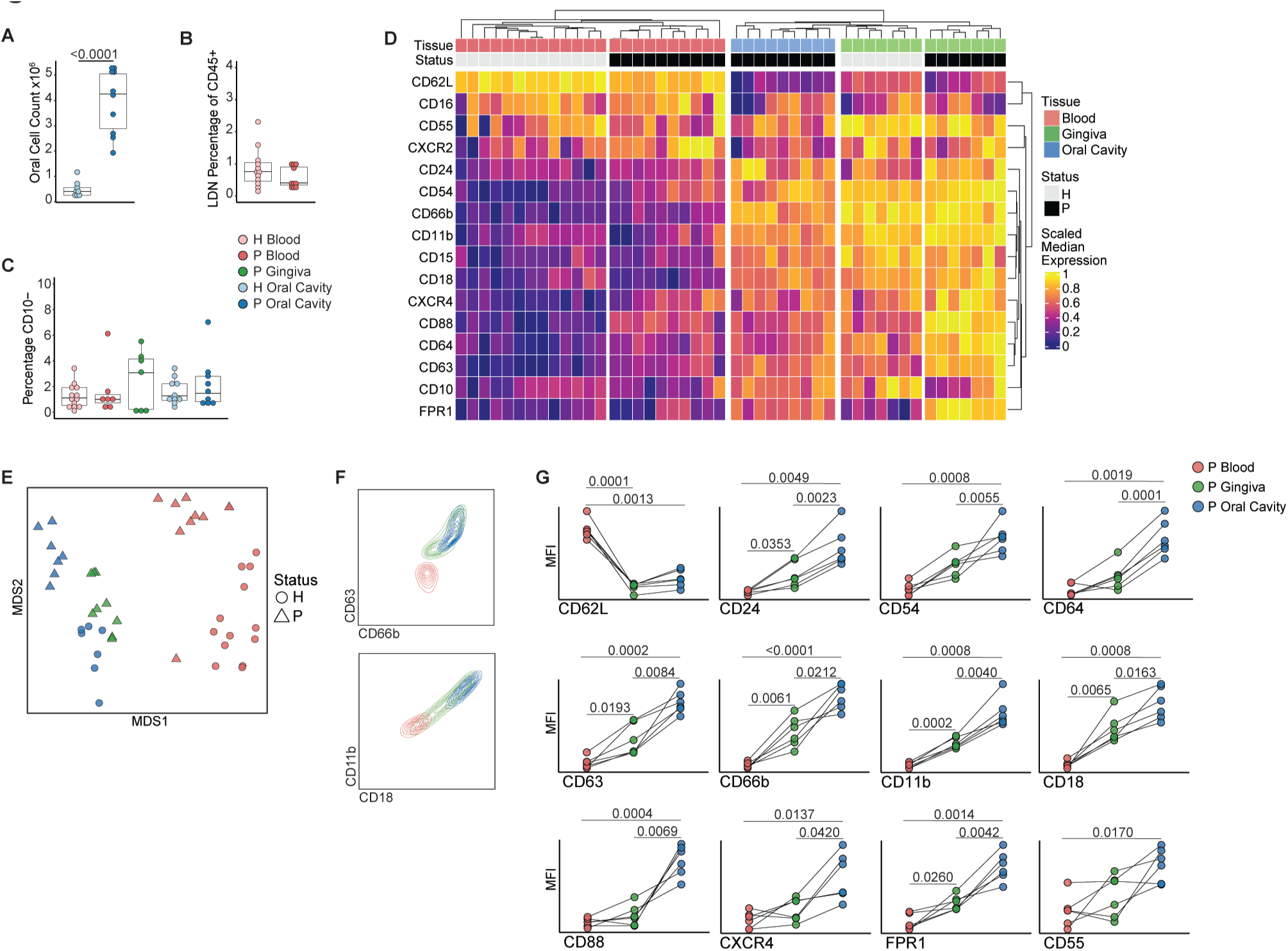
Cell surface markers reflect activation of neutrophil effector functions in the oral environment. A) Total cell counts from oral cavity sampling (n=18). B) Low density neutrophils (LDN) as percentage of the CD45+ peripheral blood mononuclear fraction (n=22). C) Percentage of CD10– neutrophils (n= 25 patients, N=46 samples). D) Heatmap of median marker expression scaled by marker across all samples. E) Multidimensional scaling (MDS) plot of H (circle) or P (triangle) patient samples from blood (red), gingiva (green), and oral cavity (blue). F) Representative contour plots from spectral flow cytometry analysis of paired P patient samples from blood (red), gingiva (green), and oral cavity (blue). G) Median fluorescent intensity (MFI) values for paired P patient samples (n=6). Data in G analyzed by repeated measures one-way ANOVA with Tukey correction for multiple comparisons.

The expression patterns of 16 established protein markers clearly separated blood neutrophils from oral tissue-derived neutrophils (**Figure 2D-E, Figure S2B**). Charting neutrophils across paired periodontitis patient samples showed a consistent reduction of CD62L expression upon entry into gingiva, while indicators of activation (CD24, CD54, CD64), degranulation (CD63, CD66b), and adhesion (CD11b, CD18) increased from blood to oral cavity with intermediate or bimodal expression in the gingiva (**Figure 2F-G**). Similarly, chemokine receptors FPR1, CD88 (C5AR1), and CXCR4 showed the highest expression in the oral cavity across subjects, suggesting a graded neutrophil response from recruitment into gingiva followed by migration into the oral cavity (**Figure 2G**). Healthy neutrophils similarly displayed a consistent decrease in CD62L and elevation in activation, degranulation, and adhesion markers expression between blood and oral cavity, (**Figure S2G**), suggesting an overall activation of neutrophils and their effector functions within the oral environment.

### Neutrophil transcriptomes define an oral tissue specification program

To further define neutrophil states and their inferred functionality within the oral mucosal environment, we evaluated neutrophil transcriptional profiles with scRNA-seq. For gingiva, we first addressed prior limitations in transcriptional profiling of human tissue neutrophils(15) by employing an alternative enzymatic digestion process coupled with a microwell-based scRNA-seq platform (BD Rhapsody)(39, 40). This approach resulted in a striking increase in the proportion of neutrophils isolated from gingiva (**Figure 3A**) and yielded a large number of high-quality gingival neutrophil transcriptomes from all subjects (**Figure 3B-E**). We supplemented this data with scRNA-seq of whole blood and unsorted oral cavity rinses, minimizing sample handling to preserve native neutrophil states.

**Figure 3.**
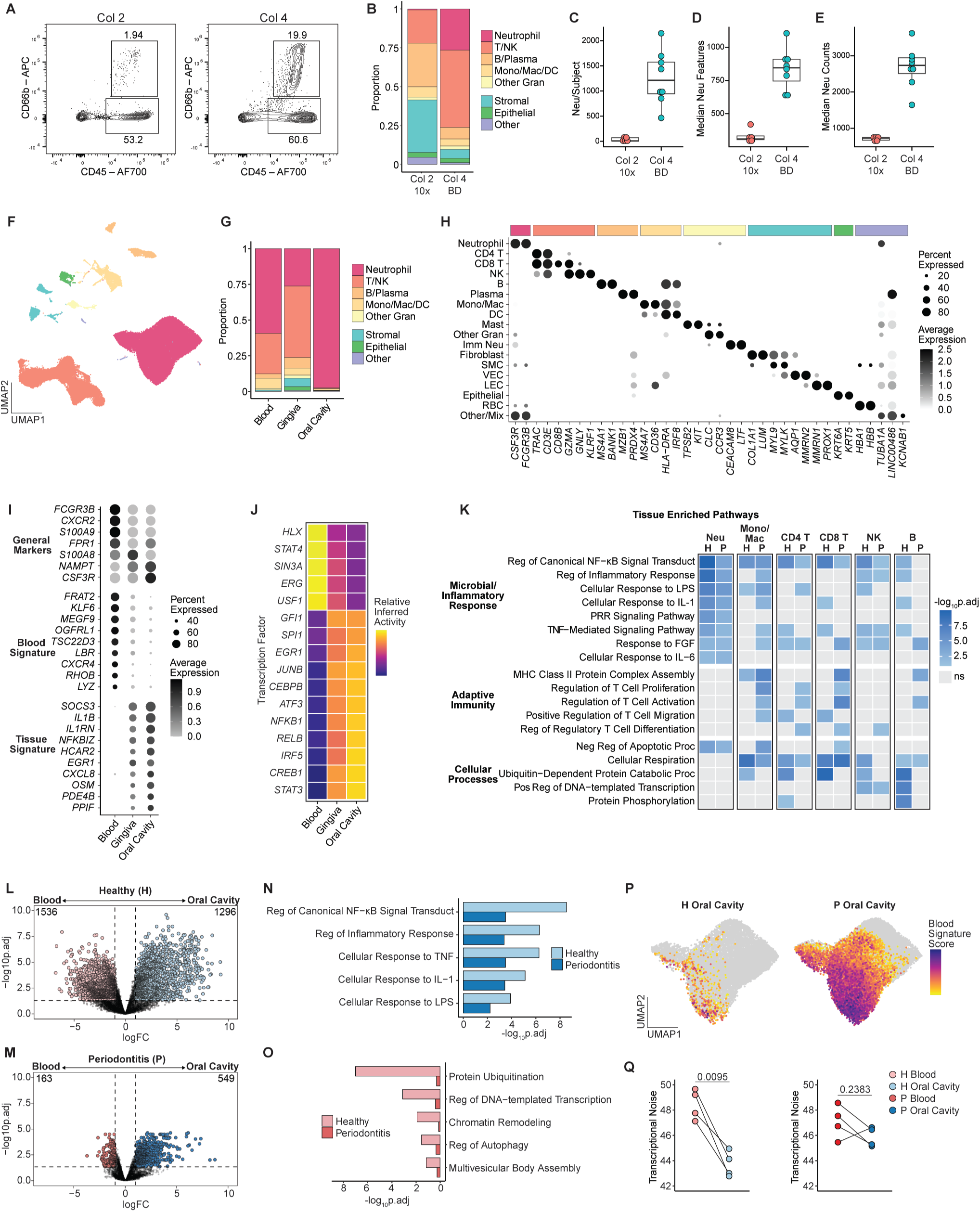
Neutrophil transcriptomes define an oral tissue specification program. A) Neutrophil recovery from gingiva digested with either collagenase 2 (Col 2) or collagenase 4 (Col 4). B) Proportion of annotated cell types in P gingiva digested with Col 2 and captured with the 10x platform (Col 2, 10x) (15) and Col 4 captured with BD platform (Col 4, BD). NK: Natural Killer, Mono: monocyte, Mac: macrophage, DC: dendritic cell, Gran: granulocyte. C-E) Quality metrics for neutrophils in respective datasets (n=5, n=8). F) Uniform manifold approximation and projection (UMAP) of combined cells from blood, gingiva, and oral cavity. G) Proportion of cell types across compartments. H) Dot plot of cell types and representative gene markers. Imm Neu: immature neutrophil, LEC: lymphatic endothelial cell, RBC: red blood cell. I) Markers distinguishing neutrophils from other cell types (General Markers), or for neutrophils in blood (Blood Signature) from gingiva and oral cavity (Tissue Signature). J) Heatmap of relative inferred transcription factor activity by compartment. K) Gene Ontology – Biologic Process (GO–BP) terms upregulated in tissue (gingiva and oral cavity) versus blood in either H or P conditions for the specified cell types. Reg: regulation, LPS: lipopolysaccharide, PRR: pattern recognition receptor, Neg: negative, Proc: process. L and M) Volcano plot showing differentially expressed genes (DEGs) for blood versus oral cavity neutrophils in either H (L) or P (M). Numbers in upper corners are the number of DEGs. N and O) GO–BP pathways upregulated (N) or downregulated (O) in oral cavity neutrophils (L) and (M). P) Visualization of Blood Signature gene score in H and P oral cavity neutrophils. Q) Transcriptional noise in paired patient samples with statistical analysis performed using paired t tests.

Analysis of 114,835 cells identified neutrophils as a prominent cellular component of each compartment, comprising more than half of blood cells, roughly 25% of gingival cells, and the vast majority (97%) of cells isolated from the oral cavity (**Figure 3F-H, Figure S3A-C**). Major cell subsets were defined based on lineage defining markers, with neutrophils expressing prototypic markers *CSF3R* and *FCGR3B* and a small number of immature neutrophils expressing neutrophil granule genes *CECAM8* and *LTF* in blood and gingiva (**Figure 3H, Figure S3**).

Further interrogation of neutrophil transcriptomes showed consistent expression of canonical neutrophil genes across tissue compartments (“General Markers” *FCGR3B, CXCR2, S100A9, FPR1, CSF3R,* and *NAMPT)* which was overlaid with distinct gene signatures based on either blood or tissue localization (**Figure 3J**). Specifically, the transition from blood to gingiva was marked by the reduction in expression of one gene set, which we termed the “Blood Signature” (*FRAT2, MEGF9, TSC22D3*), and the emergence of a “Tissue Signature” enriched in genes indicating cytokine (*SOCS3, IL1B, IL1RN, CLXCL8, OSM)* and metabolic responsiveness (*HCAR2, PDE4B, PPIF*) which was further amplified in the oral cavity (**Figure 3J**). Underlying this transition was a marked shift in inferred transcription factor (TF) activity from blood enriched TFs (*HLX, STAT4, SIN3A, ERG, USF1*), with predicted repression of Tissue Signature genes *EGR1, SOCS3*, *CXCL8* and *IL1B* (**Figure 3K, Figure S3D**), to TF programs in tissue associated with terminal neutrophil maturation (*GFI1*, *SPI1* (*PU.1*)), mobilization (*CEBPB*), and response to inflammatory stimuli (*NFKB1, RELB, JUNB, CREB1, ATF3, IRF5, STAT3*)(41, 42) (**Figure 3K**).

We next integrated this newly acquired dataset with our prior atlas(15) to compare pathways differentially enriched in oral tissue (gingiva and oral cavity) versus blood across neutrophils and other immune cells in either healthy (H) or periodontitis (P) states (**Figure 3I**). Monocyte/macrophages, CD4 T, CD8 T, and B cells in tissue showed an enrichment in inflammatory responses and T cell regulation pathways primarily in disease, consistent with their unique tissue functionalities. Oral neutrophils displayed a unique tissue signature, strongly upregulating pathways reflecting responses to cytokines and microbes and regulation of inflammation in both healthy and disease states.

To distinguish differential gene expression between blood and oral neutrophils, we performed pseudobulk comparisons in paired patient samples. Surprisingly, the number of differentially expressed genes distinguishing oral neutrophils was nearly 10-fold greater in healthy subjects compared to periodontitis (**Figure 3L-M**). Comparable inflammatory and cytokine response pathways were elevated in both health and disease, similar to single-cell level analyses (**Figure 3I**), but the downregulation of general transcriptional and protein regulation pathways was largely dampened in periodontitis (**Figure 3O**).

We reasoned that fewer genes may be differentially expressed between compartments in periodontitis due to retention of blood neutrophil features in the oral cavity. Accordingly, oral neutrophils scored with Blood Signature genes displayed a strong expression across the majority of neutrophils in periodontitis and little expression of this signature in health (**Figure 3P**). We also compared transcriptional noise, a measure of the variability in gene expression across cells(43), between subjects’ blood and oral neutrophils. In health, neutrophil transcriptional noise consistently dropped from blood to oral cavity, suggesting a reduction in stochastic neutrophil gene expression in healthy tissue, while the reduction in transcriptional noise between compartments in disease was diminished (**Figure 3Q**).

These results reveal clear neutrophil specification within the oral environment. This process was well defined in mucosal health where oral neutrophils showed distinct features from blood, while in periodontitis, the acquisition of tissue-associated features was diluted by persistent blood-associated gene programs.

### Oral specification results in distinct neutrophil transcriptional states

To further define neutrophil transcriptional states and identify possible subsets, we analyzed the combined neutrophil scRNA-seq dataset and annotated 12 clusters with unique transcriptional signatures and differential distribution in blood and oral compartments (**Figure 4A-B, Figure S4A**). Neutrophil clusters were broadly categorized into 5 groups defined by expression of either Blood Signature genes or interferon (IFN) stimulated, inflammatory, and/or regulatory gene programs (**Figure 4C**). Blood-derived neutrophils were primarily annotated in Blood Signature (clusters N1-N3) and IFN-stimulated gene groups (cluster N11-N12), while a large proportion of oral tissue neutrophils expressed signatures associated with rapid inflammatory responses, defined by immediate early genes *JUNB, IER2, EGR1* (N5-N6), or immunoregulatory programs (*TNFIAP3, PLAUR, ICAM1,* N7-N9) (**Figure 4C-E**).

**Figure 4.**
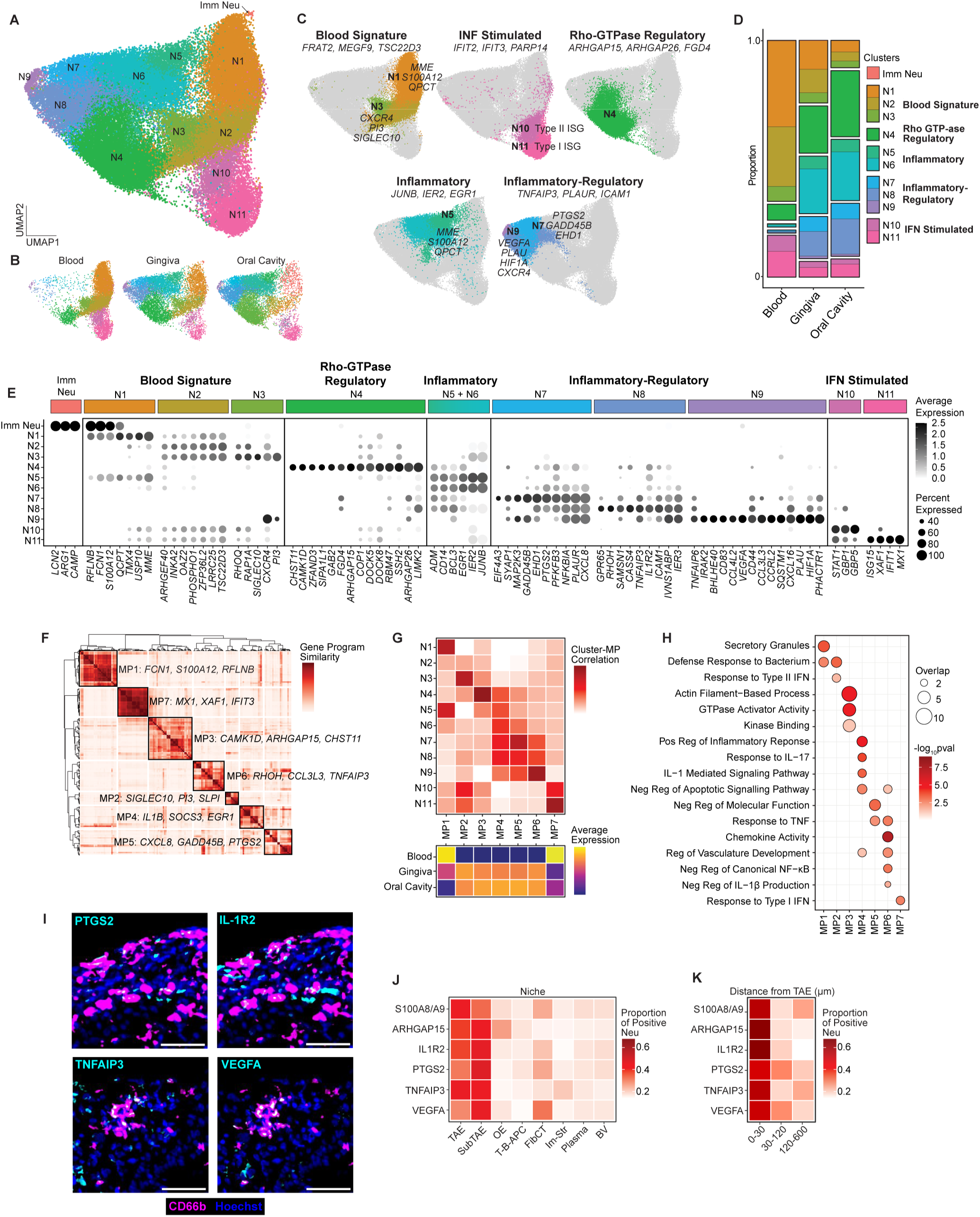
Tissue specification of oral neutrophils results in diverse transcriptional states. A) UMAP of neutrophil clusters. B) UMAP of neutrophil clusters by compartment. C) Cluster groups with associated markers and select clusters. D) Distribution of clusters across compartments. E) Top cluster markers. F) Heatmap of gene program similarities with gene metaprograms (MPs) denoted by black borders. The top 3 genes for each MP are shown. G) Heatmap showing correlation between MPs and clusters (top) and relative expression of MP genes by compartment (bottom). H) GO programs overrepresented in each MP. IFN: interferon. I) Representative images of specified neutrophil markers in P gingiva (neutrophil: magenta, nuclei: blue, marker: cyan). Scale bars are 50 μm. J and K) Distribution of neutrophils expressing each marker across niches (J) or within the given distance from the TAE (K).

Immature neutrophils (Imm Neu) were included during dimensionality reduction and clustering to infer the relative maturation status of neutrophil clusters (**Figure 4A, C, Figure S4B-C**). Two clusters of “early mature” neutrophils expressed immature neutrophil genes *FCN1* and *S100A12* and maturation marker *MME* (CD10) and either Blood (N1) or Tissue Signature (N5) genes. Two *CXCR4+, PI3+* clusters were also annotated, bearing resemblance to long-lived or aged neutrophils previously identified in blood (N3)(44) or tissue (N9)(12, 45) (**Figure 4E, C, Figure S4B-C**), with N9 neutrophils expressing additional regulatory (*TNFAIP6, CD44, CD83*), chemokine (*CCL3, CCL3L3, CCL4, CCL4L2*), and hypoxia/angiogenesis-related genes (*HIF1A* and *VEGFA*) (**Figure 4E**).

We employed an orthogonal approach, non-negative matrix factorization, to further characterize neutrophil states and identified 7 distinct neutrophil gene meta-programs (MPs), 6 of which showed high similarity to specific neutrophil clusters (**Figure 4F-G**). Two MPs showed relative enrichment in blood neutrophils: MP1, consisting primarily of genes for neutrophil granule proteins (*FCN1*, *S100A12, PGLYRP1, MMP9*) which matched the signatures of early mature N1 and inflammatory early mature N5 clusters, and MP7, with a strong type 1 IFN-stimulated signature, similar to cluster N11. MPs enriched in gingival and oral cavity neutrophils (MP2 – MP6) showed a wide range of functional gene programs including antimicrobial responses (MP2) and either positive (MP4) or negative regulation of inflammation (MP5 and MP6) (**Figure 4H**). A distinct gene metaprogram, MP3, was highly correlated with the N4 neutrophil gene signature and was uniquely enriched in genes related to GTPase and cytoskeleton regulation (**Figure 4E-H**). Overall, these tissue-associated transcriptional states infer the presence of neutrophil subsets with diverse functional programs at the oral mucosal barrier.

Finally, we evaluated gingiva for neutrophil protein expression of markers with predicted tissue enrichment, TNFAIP3 (Inflammatory Regulatory, N7-N9), PTGS2 (N7), IL-1R2 (N8) and VEGFA (N9), and identified a portion of positive neutrophils across healthy and diseased tissue (**Figure 4I, Figure S4D**). Subsets of neutrophils also expressed N4 marker ARHGAP15 and the S100A8/S100A9 heterodimer (**Figure S4E**), whose individual elements were ubiquitously expressed at the RNA level (**Figure 3I**). Protein profiles were driven by tissue regionalization, with neutrophils expressing tissue specifications markers concentrated in TAE and sub-TAE niches and in areas proximal (0-30 µm distance) to the TAE (**Figure 4J-K**).

Collectively, these findings demonstrate that oral neutrophils comprise diverse subsets with antimicrobial, pro-inflammatory, and immunoregulatory signatures. The gingiva represents the probable site where oral neutrophils acquire these programs, indicating that this barrier is a site of broad neutrophil programming.

### Inflammation disrupts tissue-specific signatures of oral neutrophils

We next evaluated the impact of disease on the distribution of oral neutrophil subsets. Intriguingly, immunoregulatory clusters N8 and N9 were enriched in health (**Figure 5A**) while clusters present in the blood (N1-N4) were overrepresented in disease, consistent with pseudobulk analysis (**Figure 3P**). To gain additional insights into oral inflammatory programing, we applied previously identified gene signatures to this data (**Figure 5B**). A core inflammatory neutrophil signature constructed from mouse and human studies(46) showed the highest expression in oral neutrophils in periodontitis. Signatures associated with classic neutrophil responses and functions (LPS stimulation, Glycolysis, Phagocytosis, and Apoptosis) also delineated healthy from periodontitis oral neutrophils, while a NETosis-related signature(47) was maximally expressed in neutrophils exiting from inflamed oral mucosa (**Figure 5b**).

**Figure 5.**
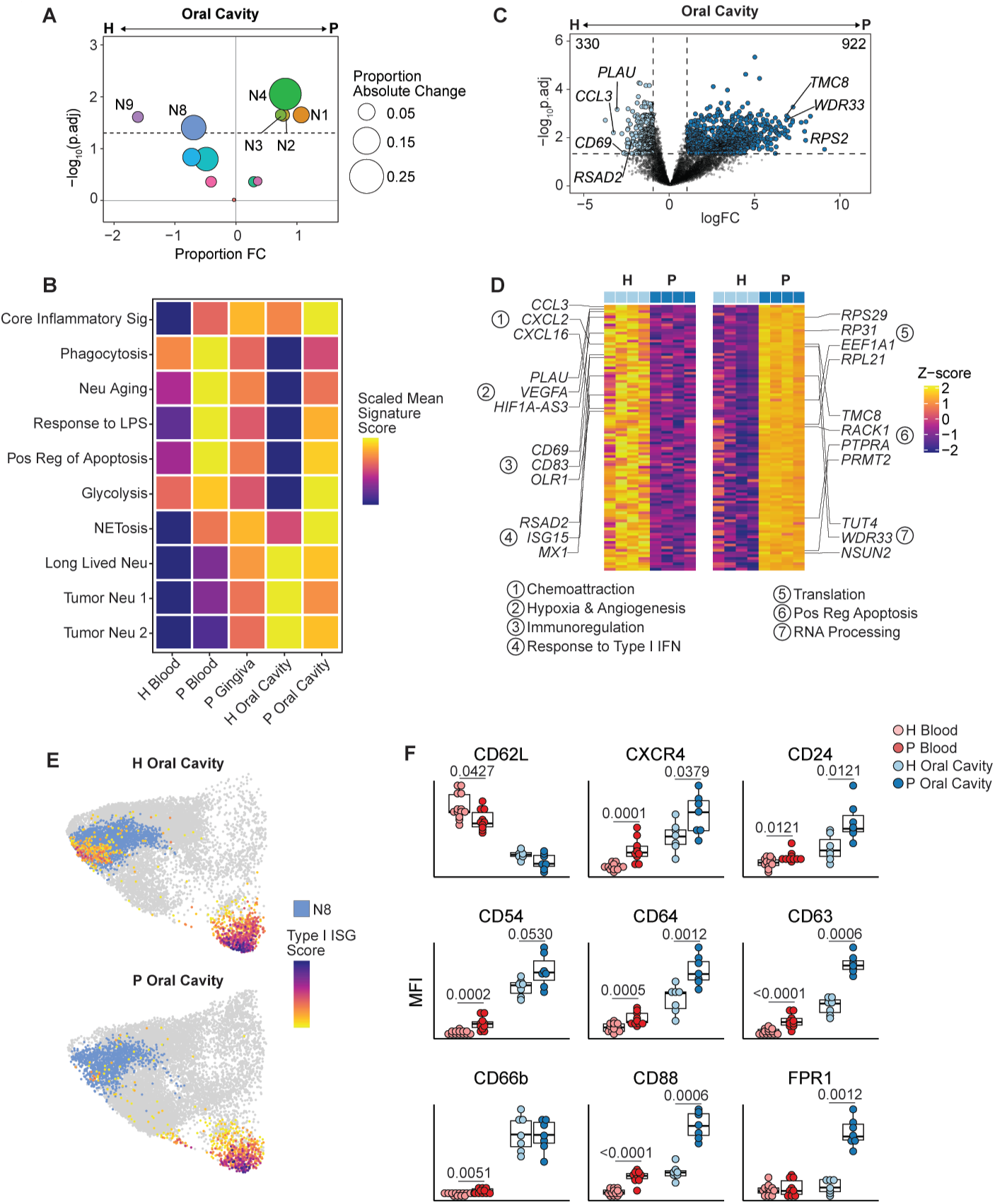
Inflammation disrupts tissue-specific signatures of oral neutrophils. A) Differences in cluster proportions between H and P oral cavity neutrophils. Symbol placement on x-axis indicates fold change (FC) in proportion and size of the symbol indicates absolute change in proportion. B) Scaled mean gene signature scores across H and P tissue compartments. C) Volcano plot of DEGs between H and P oral cavity neutrophils. Numbers in upper corners are the number of differentially expressed genes (DEGs). D) Heat map of top 100 DEGs from (C) with select genes and inferred functions from pseudobulk analysis. E) UMAP of H or P oral cavity neutrophils displaying Type I interferon-stimulated gene (ISG) score and N8 cluster annotations. F) Spectral flow cytometry marker expression in H and P blood and oral cavity neutrophils (n=25 patients, 37 samples). Data in F analyzed with individual t tests for blood or oral cavity comparisons.

At the patient level, highly upregulated genes in periodontitis oral neutrophils were tied to mRNA translation, RNA processing, and positive regulation of apoptosis pathways (**Figure 5C-D**). In contrast, healthy oral neutrophils were enriched in genes related to chemotaxis (*CCL3, CCRL2*), immunoregulation (*CREM*, *CD44, CD69, OLR1*) and angiogenesis (*PLAU, VEGFA, HIF1A-AS3*) and signatures derived from neutrophils with extended *ex vivo* lifespan (Long Lived Neu)(48) or found in tumor environments (Tumor Neu 1(40) and 2(12)) (**Figure 5B-D**). Healthy oral neutrophils further expressed higher levels of Type I interferon-stimulated genes (ISGs) (*RSAD2, ISG15, MX1*), which have been tied to neutrophil-mediated immune responses in cancer(49, 50), and displayed a distinct co-expression of the ISG signature within the inflammatory-regulatory N8 cluster (**Figure 5C, E**).

We extended our evaluation of mucosal inflammation to neutrophil surface markers. Oral neutrophils in periodontitis generally expressed higher levels of markers for activation (CD24, CD54, CD64) compared to healthy subjects. CD63, a marker of degranulation of azurophilic granules and chemokine receptors CD88 and FPR1 were strongly and consistently upregulated in disease (**Figure 5F**).

Together, these data reflect that neutrophil programing by the oral mucosa diverges between oral health and disease. Specifically, immunoregulatory and immunosuppressive neutrophils programs evident in healthy tissue were diluted by persistent blood-like subsets and enhanced inflammatory effector phenotypes in the setting of chronic inflammation.

### Oral Inflammation is imprinted on peripheral blood neutrophils

Finally, we inquired whether the effects of mucosal inflammatory disease, such as periodontitis, extend to blood neutrophil states. Gene signature scoring suggested an enrichment of inflammatory and effector programs in periodontitis versus healthy blood neutrophils (**Figure 5B**). Furthermore, subtle but significant shifts in surface marker profiles were evident in periodontitis subjects’ blood, including a reduction in CD62L signal and an increase in CXCR4 expression, a phenotype associated with neutrophil aging or activation. Similarly, markers of activation (CD24, CD54, CD64), degranulation (CD63, CD66b), and chemoattraction (CD88) in the blood were impacted by disease status, albeit to a lower degree than tissue localization (**Figure 5F**).

At the transcriptional level, blood neutrophils in periodontitis displayed increased expression of genes indicative of mRNA translation (*EEF1A1*, *RPL31, RPL15, RPS2*), and cytokine signaling and inflammatory responses (*IL6ST, NFKB1, SOCS3, IL4R, ICAM1, IL1R1)* (**Figure 6A-B**). Of note, these markers were uniformly expressed in oral neutrophils from healthy and diseased subjects, suggesting that acquisition of an activated transcriptomic state defining oral neutrophils is initiated prior to tissue entry in inflammatory disease (**Figure S5A-B**).

**Figure 6.**
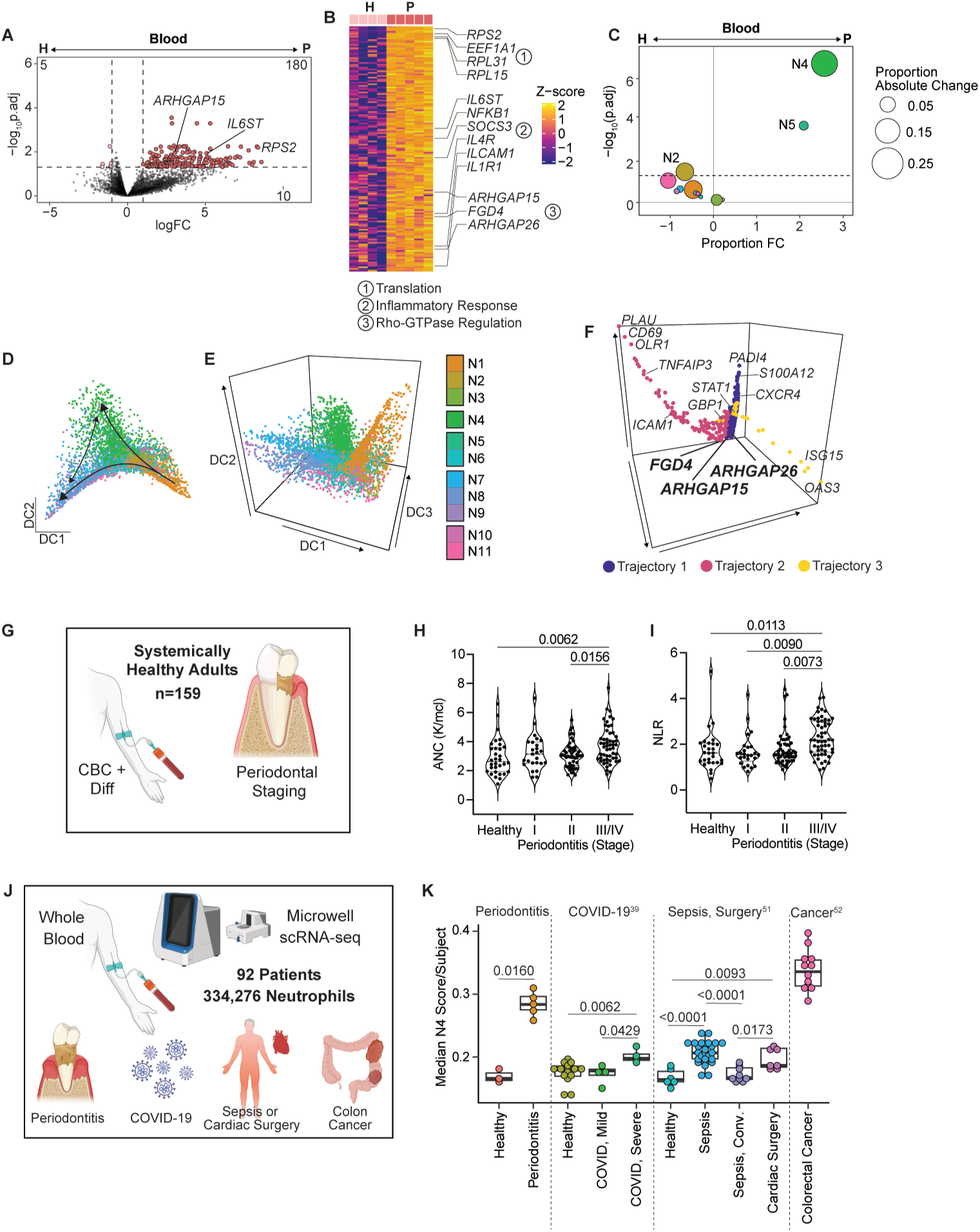
Oral Inflammation is imprinted on peripheral blood neutrophils. A) Volcano plot of DEGs between H and P blood neutrophils. B) Heatmap of top 150 DEGs from (A) with select genes and inferred functions marked. C) Differences in cluster proportion between H and P blood neutrophils. D and E) Diffusion maps of neutrophil clusters. F) Visualization of neutrophil gene trajectory with key genes marked and N4 markers in bold. G) Subjects contributing blood and receiving oral examinations. H and I) Absolute neutrophil counts (ANC) (H) and neutrophil-lymphocyte ratios (NLR) (I) in patients by periodontal diagnosis. J) Summary of studies with whole blood neutrophil scRNA-seq using BD Rhapsody included for analysis. K) Median N4 gene score in peripheral blood neutrophils by subject (symbol) across all included studies. Data in H-I analyzed via one-way ANOVA with Tukey correction for multiple comparisons. Wilcoxon-signed rank test was used in K for comparisons within individual studies with Benjamini-Hochburg adjustment used for multiple comparisons.

We additionally observed a significant upregulation of N4 signature genes *ARHGAP15, ARHGAP26,* and *FGD4* in periodontitis (**Figure 6A-B**). The N4 cluster, enriched in transcriptional programs for Rho-GTPase and actin cytoskeleton regulation, was dramatically expanded in periodontitis blood (**Figure 6C**). We examined the developmental trajectory of this cluster in the context of an inflammatory setting such as periodontitis. Plotting neutrophils with diffusion mapping suggested that N4 neutrophils could represent either the terminus of a distinct trajectory (e.g., in the blood, **Figure S4C**) or an intermediate state prior to the acquisition of an inflammatory-regulatory signature in tissue (**Figure 6D-E**). In support, inferring the sequential expression of neutrophil genes placed N4 markers *ARHGAP15, ARHGAP26,* and *CAMK1D* at the end of one trajectory indicative of neutrophil maturation and aging (Trajectory 1) and adjacent to the beginning of another pathway that terminated with expression of N9 genes *OLR1, CD69,* and *PLAU* (Trajectory 2) (**Figure 6F, Figure S5C-D**). Genes found either at the terminus of Trajectory 1 or the initiation of Trajectory 2 were highly elevated in both blood and oral neutrophils of subjects with periodontitis (**Figure S5D**), further suggesting oral mucosal inflammation induces distinct neutrophil states in peripheral blood that are maintained in the oral mucosa.

The clear impact of oral mucosal inflammation on circulating neutrophils prompted us to retrospectively investigate available blood laboratory data (complete blood counts with differential) in our larger cohort. We stratified 159 systemically healthy subjects by oral health status into periodontally healthy or Stage I (mild), Stage II (moderate), or Stage III/IV (severe) periodontitis groups. Both absolute neutrophil count (ANC) and neutrophil-lymphocyte ratio (NLR) were significantly elevated in severe periodontitis, despite falling within normal clinical ranges (**Figure 6G-H**). Further, periodontitis disease severity was positively correlated with both ANC and NLR, which remained significant when adjusting for age, sex, and ethnicity (**Figure S5E-H**).

Finally, these intriguing findings, revealing systemic imprinting of mucosal inflammation and the emergence of a distinct neutrophil transcriptional state, prompted us to inquire whether the peripheral N4 gene signature was a unique feature of periodontal disease or a signature of systemic neutrophil conditioning secondary to tissue and/or systemic inflammation. We queried available scRNA-seq datasets acquired with the same microwell-based platform across diverse inflammatory diseases. Evaluation of 334,276 blood neutrophils across 92 subjects showed low levels of N4 gene signature in healthy subjects while circulating neutrophils from acute or chronic inflammatory conditions such as severe COVID-19(39), sepsis and cardiac surgery(51), or colorectal cancer(52) demonstrated a significant enrichment in the N4 genes (**Figure 6I**) as well as signatures for both Rho-GTPase inhibitors (GTPase activators, GAPs) and activators (guanine nucleotide exchange factors, GEFs) (**Figure S5I-J**).

These findings, together, demonstrate that oral mucosal inflammation imprints on circulating neutrophils at the transcriptomic and proteomic level. This imprinting is marked, in part, by a molecular signature of Rho-GTPase and cytoskeletal regulation which emerges as a shared transcriptional state across diverse human inflammatory conditions.

## Discussion

Our work defines neutrophil transcriptional programs across interconnected compartments of blood, oral mucosa (gingiva), and oral cavity and describes tissue specification programs of human neutrophils in health and inflammatory disease. Moreover, our study reveals an imprinting of neutrophil states beyond the tissue microenvironment on circulating neutrophils, with the emergence of a distinct transcriptional state that is shared across various human inflammatory conditions.

A strength of our approach is the meticulous clinical study design which defines health and oral inflammatory disease with strict criteria while minimizing confounding factors known to affect neutrophil biology. As such, our cohorts were carefully curated to include subjects with either pristine oral health or untreated severe inflammatory disease in the absence of medical co-morbidities, medications such as antibiotics and anti-inflammatory drugs, and smoking, among other stringent criteria(53). Furthermore, individuals were evaluated and sampled within the same timeframe to control for variations in circadian rhythm(11) and samples were immediately processed using methodologies optimized for neutrophils and oral tissues. Importantly, matched samples were obtained within individuals to track neutrophil specification programs across compartments. Conceivably, this approach allowed us to track the path of neutrophils from the circulation, into the gingiva, and out to the oral cavity, which is well established as a pathway of continuous neutrophil migration in humans(54).

We visualize neutrophils in the gingiva and observe that this highly mobile immune cell type forms an integral part of the tooth-facing barrier. Neutrophils are found consistently within the gingival TAE in health and greatly expand in numbers in the setting of periodontitis(55, 56), extending into adjacent sub-epithelial tissue(21). The TAE itself expresses diverse neutrophil chemotactic signals in health which are markedly increased in disease(21, 57), indicating that the TAE is a driving force behind oral neutrophil recruitment. In other barriers such as the intestine, intraepithelial neutrophils are an indicator of inflammatory disease activity while the presence of sub-epithelial neutrophils is more characteristic of homeostatic immunity(58, 59). Accordingly, epithelial damage by migrating neutrophils and subsequent loss of barrier function has been identified in diseased lung, urinary tract, and intestines(60, 61). In contrast, the gingival TAE is characterized by wide intraepithelial spaces with few tight junctions which allows neutrophil migration with minimal epithelial damage but also exposes this interface to microbial insult and injury. The band-like organization of neutrophil, along with the distinctive presence of NETs within and below the TAE, further suggests that neutrophils physically secure this permeable barrier in health and are crucial in periodontitis where the TAE increases in permeability and/or ulcerates and microbial translocation increases(62, 63).

Beyond forming a structural barrier through NETs, neutrophils conceivably mediate immune surveillance and active antimicrobial defense at the TAE and participate in ongoing wound healing events that are essential in this continuously microbially triggered and injured interphase(64). However, whether neutrophils acquire a distinct transcriptional program within mucosal tissues and how their functionality is altered in health versus inflammation is incompletely understood across mucosal sites in humans.

We detect a clear oral tissue specification program in human neutrophils that is distinct from circulating blood neutrophils within the same individuals. This specification reflects the emergence of cellular states with different inferred functionalities. At the protein level, oral neutrophils upregulate markers consistent with cell activation and induction of effector functions, such as degranulation and NETosis, consistent with the homeostatic inflammation and local antimicrobial defense that defines this barrier, even in the absence of clinical disease(21). At the transcriptional level, neutrophil subsets that appear in the mucosa encode both inflammatory and immunoregulatory gene programs. Tissue inflammatory programs are characterized by the expression of rapidly acquired inflammatory genes (*JUNB, IER2, EGR1)*(65), as well as broader activation states (*IL1B, NFKBIA, CXCL8*). Intriguingly, neutrophil subsets with clear immunoregulatory profiles emerge as a hallmark of oral neutrophil specification, particularly in health. As such health-enriched neutrophil subsets express genes associated with the dampening of cytokine responses and NF-κB signaling (*IL1R2, TNFAIP3, TNFAIP6*) and upregulate programs previously associated with tumor neutrophils (*OLR1, PLAU, VEGFA, HIF1A*, etc.)(12, 40) and cancer immunotherapy (Type 1 ISGs)(49, 50). These findings lend support to the notion that loss of neutrophils and their regulatory roles within barrier tissues, and particularly within the oral environment, severely disrupts homeostatic immunity(25, 66). It is recognized that neutrophils can indirectly modulate inflammatory signaling through their apoptosis to regulate the IL-23–IL-17 axis(67). However, whether particular neutrophils subsets play active, immunoregulatory roles within barrier tissues has not been detailed to date. Furthermore, our data suggests active regulatory effects, beyond indirect regulatory roles, for certain neutrophil subsets that appear native to the mucosal environment. These activities may range from acting as cytokine sinks through IL-1R2 to secreting immunomodulatory mediators (COX-2 (*PTSG2),* TSG-6 (*TNFAIP6*)) and regulating their own cytokine response signaling (A20 (*TNFAIP3*)), suggesting these cells act as active orchestrators of immunoregulation at the healthy oral barrier.

In disease, neutrophil immunoregulatory programs are diluted by the persistence of blood-like signatures within the mucosa. We hypothesize that elevated inflammation and microbial triggering in periodontitis amplify neutrophil chemotactic signals in the TAE and underlying stromal cells(15, 21) which accelerates neutrophil recruitment from the blood. The existence of additional transcriptional states within the tissue, conceivably retained from blood, together with the plethora of additional stimuli within the inflamed tissues, may explain the increased transcriptional noise in periodontitis neutrophils(43).

This altered state can theoretically provide neutrophils with an increased plasticity of response within the more challenging and threatening environment of periodontitis(68, 69). We also confirm prior findings that oral neutrophil states are altered at the proteomic level in periodontitis, consistent with increased activation by inflammatory and microbial stimuli(70, 71). While this exaggerated inflammatory state is tied to antimicrobial effector functions, excessive neutrophil recruitment and activation is also linked to oral immunopathology and may contribute to disease severity in periodontitis(28, 29, 31, 72). Strikingly, oral mucosal inflammation imprints on neutrophils states beyond the oral environment. We identify a clear upregulation of neutrophil activation markers and an expansion of particular neutrophil transcriptional states in the blood. Acute and severe systemic inflammation has been typically associated with emergency granulopoiesis, manifested by an increase in neutrophil numbers and release of immature or irregularly mature neutrophil states from the bone marrow(4, 51). However, periodontitis by itself does not appear to induce a classic emergency granulopoiesis response, as increases in circulating neutrophil numbers are modest and immature and/or low-density neutrophils are not significantly increased. Yet systemic imprinting is clear in this study and further builds on prior work demonstrating persistent impacts of periodontitis on peripheral neutrophil functions like reactive oxygen species and cytokine production(73, 74). Whether this altered neutrophil state results from conditioning of circulating neutrophils, secondary to systemic release of inflammatory and microbial factors from oral tissues(75, 76), or represents altered granulopoiesis as a result of trained innate immunity at the progenitor level(35), cannot be determined in the current study.

Moreover, we document the expansion of a particular neutrophil subset in the circulation of subjects with periodontitis reflecting the regulation of Rho-GTPase activity through both activators (GEFs *FGD4, DOCK5, DOCK8,* etc.) and inhibitors (GAPs *ARHGAP15, ARHGAP26, RASA2*, etc.) of these signaling proteins. Rho-GTPases, including RAC1, RAC2, and CDC42 play important roles in cytoskeletal organization(77) and are implicated in a wide range of neutrophil functions, including migration, recruitment, degranulation, phagocytosis, NADPH oxidase activity, and NETosis(78, 79). As such *RAC2* mutations are linked to immunodeficiency syndromes affecting neutrophil motility and function(80, 81) and defects in the GEF, *DOCK2,* have been tied to impaired neutrophil actin polymerization and decreased oxidative burst activity(82, 83).

Remarkably, analysis of publicly available datasets reveals that this neutrophil transcriptional state is not unique to periodontitis but is also expanded in patients with diverse inflammatory disease states ranging from sepsis to malignancy. These data introduce the Rho-GTPase regulator-enriched neutrophil state as a shared program reflecting systemic conditioning of peripheral neutrophils by inflammatory disease. Whether this subset is endowed with unique functionality is unclear, but its existence opens avenues towards understanding global programs that underpin the crosstalk between tissue immunity and peripheral neutrophil conditioning.

Collectively, our work provides a comprehensive characterization of human neutrophil states across blood, oral mucosa, and oral cavity compartments, revealing a transcriptional diversification in the gingiva with the acquisition of inflammatory and regulatory gene programs. Chronic mucosal inflammation alters this specification and drives the expansion of a neutrophil and NET niche at the tooth-mucosal interface. Critically, our results highlight the systemic consequences of dysregulated barrier immunity, demonstrating that chronic inflammation extends mucosal conditioning of neutrophils beyond the oral cavity. These findings provide a framework for understanding how barrier tissue environments shape neutrophil biology with implications extending beyond oral health to systemic inflammatory diseases.

## Acknowledgements

This research was supported by the Intramural Research Program of the National Institutes of Health (NIH): Division of Intramural Research of the National Institute of Allergy and Infectious Diseases and the Intramural Program of the National Institute of Dental and Craniofacial Research. This research was supported, in part, by the NIDCR Combined Technical Research Core (ZIC DE000729-09) and Imaging Core (ZIC DE000750). This work utilized the computational resources of the NIH HPC Biowulf cluster (https://hpc.nih.gov). The contributions of the NIH authors are considered Works of the United States Government. The findings and conclusions presented in this paper are those of the author(s) and do not necessarily reflect the views of the NIH or the U.S. Department of Health and Human Services. We also acknowledge scRNA-seq technical and sequencing support from Michael Kelly and the staff of the National Cancer Institute, Center for Cancer Research Single Cell and Spatial Core, sequencing support from Daniel Martin and the NIDCR Genomics and Computational Biology Core, and imaging support from Andrew Doyle in the NIDCR Imaging Core. Figure illustrations were created with Biorender.com or by Ryan Kissinger in the NIAID Research and Technologies Branch.

## Author Contributions

DF and NMM designed the study and wrote the manuscript. DF performed single-cell sample collection, experimental work, and data analysis. VT performed spatial experiments and data analysis. TGW and CW assisted with experimental work. TGW, LG, and EK assisted with patient enrollment and sample collection. NMM supervised the study. All authors reviewed and edited the manuscript

## Methods

### Human subjects

Subjects were screened through an ongoing clinical protocol at the National Institutes of Health Clinical Center (ClinicalTrials.gov ID NCT01568697) and enrolled if eligible. Eligibility criteria included age ≥18 years, the presence of 20 or more natural teeth, and overall systemic health. Exclusion criteria included a prior diagnosis of diabetes or HbA1C levels >6%, any autoimmune disorder, current pregnancy or lactation, active malignancy, prior radiation to head or neck, history of HIV or hepatitis B or C infection, use of antibiotics, corticosteroids, immunosuppressants, cytokine therapies, high-dose commercial probiotics within the last 3 months, or use of tobacco products or e-cigarettes within the last 1 year. Following informed consent, enrolled subjects received a comprehensive oral examination, oral radiographs, and clinical blood laboratory testing. Subjects were diagnosed as having periodontal health, gingivitis, or periodontitis (Stage I, II, III or IV) based on clinical and radiographic findings(84). Subjects with either periodontal health or untreated, generalized (>30% of sites affected) Stage III or IV periodontitis (Grade B or C) provided additional blood, gingiva, and/or oral rinse samples.

### Sample Collection

All samples were acquired between 8 and 11 AM from subjects who had not eaten, drank, or brushed their teeth for least 90 minutes prior. Ten mL blood was collected in sodium heparin tubes and processed for either neutrophil isolation by density gradient centrifugation with Ficoll Paque Premium(85) or subjected to whole blood magnetic RBC depletion (EasySep RBC Depletion Reagent). Gingiva for single-cell suspensions was obtained from tissue discarded during surgical treatment of periodontitis subjects using a method modified from our prior study(86). Briefly, surgical discards were collected in cold RPMI media containing 2% fetal bovine serum and 0.04 mg/mL DNase 1 (Sigma). Loose and fibrous tissues were separated from each other by gentle scraping with a scalpel blade. The loose tissue fraction was minced and immediately pressed through a 70-um cell strainer with a syringe plunger. The fibrous fraction was processed to single cell suspensions as previously described but substituting collagenase 4 (1 mg/mL) for collagenase 2 and limiting incubation at 37° C to 30 minutes. Both fractions were then combined and RBCs were lysed with a hypotonic lysis solution. For scRNA-seq applications, dead cells were magnetically depleted (EasySep Dead Cell Removal Kit) to reach a viability of ≥80%. Oral rinse samples were collected by having subjects rinse twice with 10 mL of 0.9% sodium chloride for 30 seconds, with each rinse separated by a 3-minute interval. Oral rinses were immediately diluted in ice cold calcium- and magnesium-free PBS supplemented with 0.04 mg/mL DNase 1 and centrifuged, followed by filtering through 40 um and 20 um cell strainers to remove debris and deplete epithelial cells.

### Spatial Proteomics Staining and Imaging

Gingiva for spatial proteomics were obtained from healthy and periodontitis subjects as previously described(21). Briefly, biopsies from healthy sites were taken from clinical sites without visible inflammation, probing depths ≤3 mm, no bleeding on probing, and no radiographic bone loss. Periodontitis subject biopsies were taken at sites of severe disease (probing depths and clinical attachment levels ≥ 5 mm) with active inflammation. Tissue orientation was carefully recorded prior to grossing and fixation in 1% paraformaldehyde, followed by incubation in 30% sucrose and embedding in cryo-embedding medium. 5 μm sections from all tissues were placed on a single chrome alum gelatin-coated cover glass which was then attached to a 3D printed microscope stage insert containing a chamber for staining reagents. Sections were rehydrated and blocked with a solution of 0.3% Triton-X-100, 1% BSA, 1% human Fc block (BD Biosciences) and 0.02% Hoechst (Biotium) at 37 °C for 1 hour. Primary antibodies were incubated overnight at 4 °C, followed by secondary antibody incubation at 37 °C for 1 hour (**Table S17**). For biotin-labeled primary antibodies, sections were treated with endogenous streptavidin/biotin blocking solutions (Invitrogen) prior to antibody incubation.

Images were acquired on a Nikon A1R HD25 confocal microscope equipped with 488 nm, 561 nm and 647 nm laser lines and a 20x air objective (Plan Apo VC 20x DIC with 0.75 numerical aperture). Resonant scanner mode was employed with 4x line averaging, a pinhole size of 2.9 AU, and pixel dimensions of 0.62 μm x 0.62 μm with a 12-bit depth. Z-stacks containing whole sections were obtained and raw outputs were then pre-processed at NIS Elements (Nikon) with denoising, multi-tile stitching, and maximum intensity projection.

### Spatial Proteomics Image Processing and Analysis

Files were processed and registered based on nuclei channel morphology as previously described(21, 87). Registered multichannel images were processed automatically in ImageJ. First, a combination of automatic and manual removal of unregistered or detached section areas was conducted. Individual channels were then processed with median filter, background subtraction, thresholding, and small object elimination and the area below the intensity and object size threshold was eliminated from each channel. Cell segmentation was conducted with a combination of Ilastik(88), and Cellprofiler(89) using nuclei channel morphology to reduce overlaps between adjacent cells in highly cellular areas. Quantification of per channel intensities was conducted using the processed multichannel images and the segmentation outputs with MCQuant.

Clustering analyses were conducted with scimap(90). Protein marker intensity values were scaled via the rescale function using a Gaussian Mixture Model for automatic scaling. Initial coarse k-means clustering was carried out to define major cell types followed by sub-clustering within each cluster. A subset of relevant markers was used when appropriate for cell type clustering. For niche analysis, the spatial_expression tool was used to generate weighted neighborhood matrices based on the k-nearest neighbors (kNN=30), followed by determination of the k-means of the neighborhood matrices using the spatial_cluster tool. Cell types and niche clusters were manually merged when appropriate, based on shared protein marker expression and histologic context. Pathologist annotations were applied with the addROI_image tool to ensure histologically accurate clustering of epithelial cell types and niches.

To assess the spatial distribution of neutrophils relative to the TAE niche, pairwise Euclidean distances were calculated between all cells and TAE cells using the BallTree algorithm implemented in scikit-learn based on X and Y centroid coordinates. Distance calculations were performed up to a maximum threshold of 1000 pixel-units to account for variability in tissue section size. Cells were subsequently stratified into radial distance bins from the TAE. Using a similar spatial framework, neutrophil neighborhood relationships were evaluated by calculating the average distance of surrounding cell types to neutrophil clusters and the number of neighboring cells within a defined proximity radius (30-pixel units/18 μm).

NET area quantification was performed in ImageJ using CitHist and MPO channels. Briefly, each channel underwent automated preprocessing including background subtraction, median filtering, and thresholding. Segmented objects with particle size of 120-pixel units or greater (substantially exceeding the average neutrophil particle size of 20-pixel units) were retained to ensure accurate representation of extracellular NET structures. To increase specificity, only CitHist-positive objects demonstrating spatial co-localization with the MPO mask were classified as NETs. NET areas were quantified and normalized to total tissue area. NET spatial coordinates were exported for downstream analysis in Python. Euclidean distances were then calculated between all cell centroids and NET area outlines from the corresponding tissue section. These measurements were used to determine the average distance of each cell type and niche to NET regions and the number of cells located within a defined proximity radius of NET areas (30-pixel units/18 μm).

### Full Spectrum Flow Cytometry

Cell suspensions were first incubated with Zombie NIR viability dye, then rinsed, blocked with normal mouse serum, incubated with antibodies at 4° C (**Table S18**), and fixed with 4% paraformaldehyde prior to analysis on a Cytek Aurora 5 laser full spectrum flow cytometer. Outputs were unmixed with SpectroFlo software and transferred to FlowJo for gating of singlet, live, CD45+, CD66b+, CD15 high, CD16+ neutrophils. FCS files were exported to R for data visualization and analysis with CATALYST (91).

### scRNA-seq Data Acquisition

Single cell suspensions were prepared in Sample Buffer (BD) with 0.2 U/μL RNase inhibitor (Roche) and loaded onto BD Rhapsody capture cartridges with Cell Capture Beads following the manufacturer’s instructions. Multiplexing for a subset of samples was performed prior to capture using the BD Human Single Cell Multiplexing Kit. Following cell lysis, beads were isolated and reverse transcription and exonuclease treatment was performed. Libraries were prepared with BD Rhapsody reagents and checked for proper concentration and peak size distribution prior to sequencing with the NovaSeq X platform (Illumina). Fastq file filtering, alignment, doublet identification, and annotation with GRCh38 was performed on the Seven Bridges cloud platform using the BD Rhapsody Sequence Analysis Pipeline.

### scRNA-seq Data Processing and Cluster Analysis

All scRNA-seq data processing and analysis was performed in R. Counts were processed to remove ambient RNA with SoupX(92). All subjects’ counts were merged and cells with fewer than 100 or more than 6000 features and/or >25% mitochondrial reads were filtered out, with the latter threshold chosen based on prior studies showing a higher mitochondrial content in cells captured with the BD Rhapsody platform(39, 93). Log normalization, determination of variable genes (n=4000), scaling, and PCA reduction were performed with Seurat v5(94). Harmony was employed for batch correction using the individual subject sample as the covariate(95). UMAP reduction and nearest-neighbor graph construction was performed using dimensions 1:50 and clusters were created using the Louvain algorithm with a resolution of 1. The FindAllMarkers function in Seurat was employed to identify differentially expressed genes by cluster and clusters were manually annotated by referencing published single cell atlases of blood and gingiva(15, 51).

Neutrophil and immature neutrophil clusters were subset from all cells and re-processed by removing cells with >10% mitochondrial reads followed by determining variable genes (n=2000), scaling, performing PCA reduction and Harmony batch correction using the individual sample as the covariate, running UMAP reduction and nearest-neighbor graph construction with dimensions 1:30, and finding clusters using the Louvain algorithm with a resolution of 1. Clusters expressing markers defining other cell types (possible heterotypic doublets), primarily mitochondrial or ribosomal genes, or only heat shock protein genes *HSPA5* and *HSP90B1* (considered poor quality cells) were removed and processing was repeated. Broad and fine-level neutrophil clusters were determined by comparing cluster stability across a range of resolutions with clustree(96). Cluster distribution between conditions or tissues was calculated using precision weighted linear mixed models(97), aggregating cells from each subject sample and including batching and subject sex as model covariates when determining differential

cluster abundance.

### Transcription Factor Inference

Inferred transcription factor activity and targets were estimated using univariate linear models with decoupleR(98). DoRothEA databases were queried for transcription factor targets(99).

### Estimation of Transcriptional Noise

Transcriptional noise for each subject sample was estimated by calculating Euclidean distances between the mean gene expression vector of all sample neutrophils and the gene expression vector of each individual neutrophil within the sample.

### Non-negative Matrix Factorization (NMF)

NMF was performed using geneNMF(100). Variable features were calculated (n=2000) while excluding mitochondrial genes. Metaprograms (MPs) containing more than 10 programs and a silhouette >0.2 were retained. Neutrophils were scored for MP genes and cluster markers using UCell(101) to determine MP – cluster correlations. Enriched gene programs for each MP were calculated with the runGSEA function using all Gene Ontology gene sets.

### Pseudotime and Cell and Gene Trajectory Analysis

Neutrophil pseudotime was estimated using Monocle(102). Neutrophils were split by tissue and pseudotime was calculated for each tissue separately. Diffusion mapping of neutrophils was performed with Destiny(103). Neutrophil gene trajectory was calculated with GeneTrajectory(104) using genes expressed by >3% and <50% within neutrophil variable genes (n=3000). Gene embedding and trajectory was performed using K=5.

### Integration with Published Datasets

Count matrices and metadata data from the current study and a prior atlas of healthy and periodontitis gingiva(15) were merged, filtered, and integrated using Harmony with individual subject sample and the study platform (10x Chromium or BD Rhapsody) as covariates. Neutrophils were subset and re-processed to remove heterotypic doublets prior to quantifying neutrophil quality metrics. To compare blood or tissue enriched pathways in health or periodontitis, overrepresentation analysis was performed with Enrichr(105) for major cell types with adequate representation, defined as >1000 cells of each cell type present in both blood and tissue (gingiva and oral cavity combined) in both health and periodontitis.

Studies with publicly available data for scRNA-seq of whole blood acquired on the BD Rhapsody platform were processed as above (COVID-19(39): https://uni-bonn.sciebo.de/s/7ZOxo0nc448orgT?path=%2FDatasets, Sepsis and cardiac surgery(51): https://zenodo.org/records/7723202, Colon Cancer(52): https://crc.icbi.at/). The N4 blood gene signature was derived using the FindAllMarkers function for blood neutrophils from the current study and applied to all neutrophil datasets using UCell. Counts were extracted and scores were aggregated at the subject level.

### Pseudobulk Analysis

Cell counts were aggregated at the cell type and subject level and normalized, and differential gene expression was calculated using Dreamlet(106) with subject sex and within-subject batching as covariates. Overrepresentation analysis for pairwise group comparison was performed using Enrichr and Gene Ontology: Biologic Process databases. Individual Neutrophils were scored for gene signatures (**Table S13**) using UCell and the mean signature score was aggregated by subject.

## Data Availability

scRNA-seq data have been deposited in GEO under accession GSE325501.

## Supplemental Materials

Figures S1-S5

Table S1. Subject Metadata

Table S2. scRNA-seq Cell Type Markers (Figure 3H)

Table S3. Neutrophil scRNA-seq Markers by Compartment (Figure 3I)

Table S4. Immune Cell Enriched Pathways: Periodontitis Tissue vs. Blood (Figure 3K)

Table S5. Immune Cell Enriched Pathways: Healthy Tissue vs. Blood (Figure 3K)

Table S6. Healthy Oral Cavity vs. Blood Differentially Expressed Genes (Figure 3L)

Table S7. Periodontitis Oral Cavity vs. Blood Differentially Expressed Genes (Figure 3M)

Table S8: Oral Cavity vs. Blood Enriched Pathways (Figure 3N)

Table S9. Blood vs. Oral Cavity Enriched Pathways (Figure 3O)

Table S10. Neutrophil Cluster Markers (Figure 4A-E)

Table S11. Neutrophil MP Markers (Figure 4F)

Table S12. Neutrophil MP Pathways (Figure 4H)

Table S13. Gene Signatures for Scoring

Table S14. Oral Cavity Healthy vs. Periodontitis Differentially Expressed Genes (Figure 5C-D)

Table S15. Blood Healthy vs. Periodontitis Differentially Expressed Genes (Figures 6A-B)

Table S16. Neutrophil Gene Trajectories (Figure 6D-E, Figure S5C-D)

Table S17. Spatial Proteomics Antibodies

Table S18. Spectral Flow Cytometry Antibodies

## Supplemental Figure Legends

**Figure S1.**
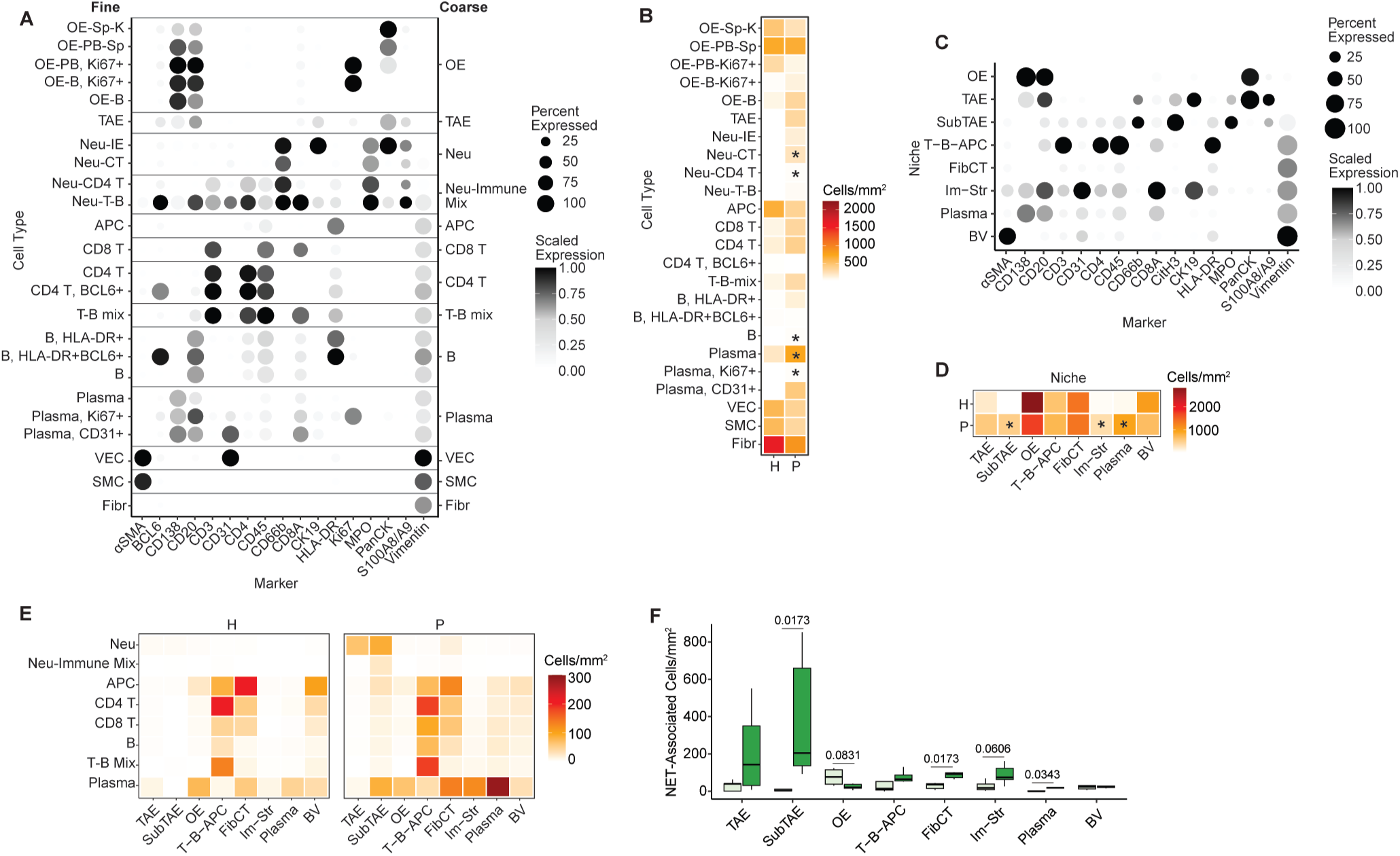
Corresponding to Figure 1. A) Dot plot of markers and fine (left) or coarse (right) cell type annotation for spatial proteomics. OE: oral epithelium, SP: spinous, K: keratin, PB: parabasal, TAE: tooth-associated epithelium, Neu: Neutrophil, IE: intra-epithelial, CT: connective tissue, APC: non-B antigen presenting cell, VEC: vascular endothelial cell, SMC: smooth muscle cell, Fibr: fibroblast. B) Mean number of cells per area (mm^2^) per cell type in healthy (H) and periodontitis (P) subjects. C) Dot plot of cell marker expression within indicated tissue niches. FibCT: fibrous connective tissue, Im-Str: immune-stromal, BV: blood vessel. D) Mean cell density within each niche in H and P subjects. E) Density of indicated immune cell types across niches within H and P gingiva. F) NET-associated cells per niche, normalized by tissue area, in H and P subjects. N=11. Wilcoxon-signed rank test was used for B, D, and F with Benjamini-Hochburg adjustment for multiple comparisons. Asterisks in B and D represent adjusted p-values <0.05 and lines with numbers in H are adjusted p-values.

**Figure S2.**
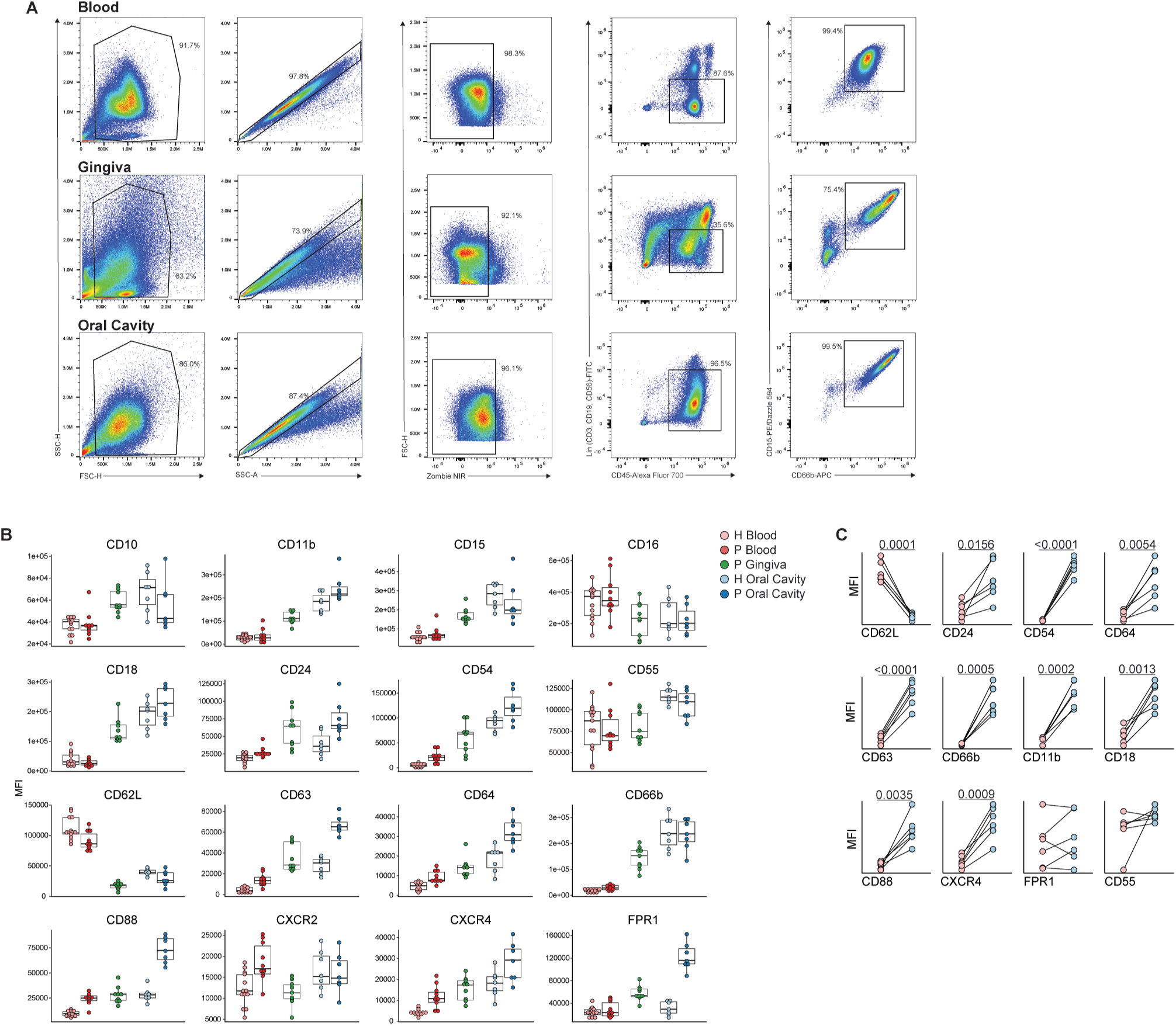
Corresponding to Figure 2, Spectral flow cytometry. B) Gating strategy for identification of neutrophils. B) Per patient MFI values for indicated markers on neutrophils from H and P blood and oral cavity samples corresponding to Figure 2D. C) MFI values for paired H samples from blood and oral cavity (n=6). Data analyzed by paired t test.

**Figure S3.**
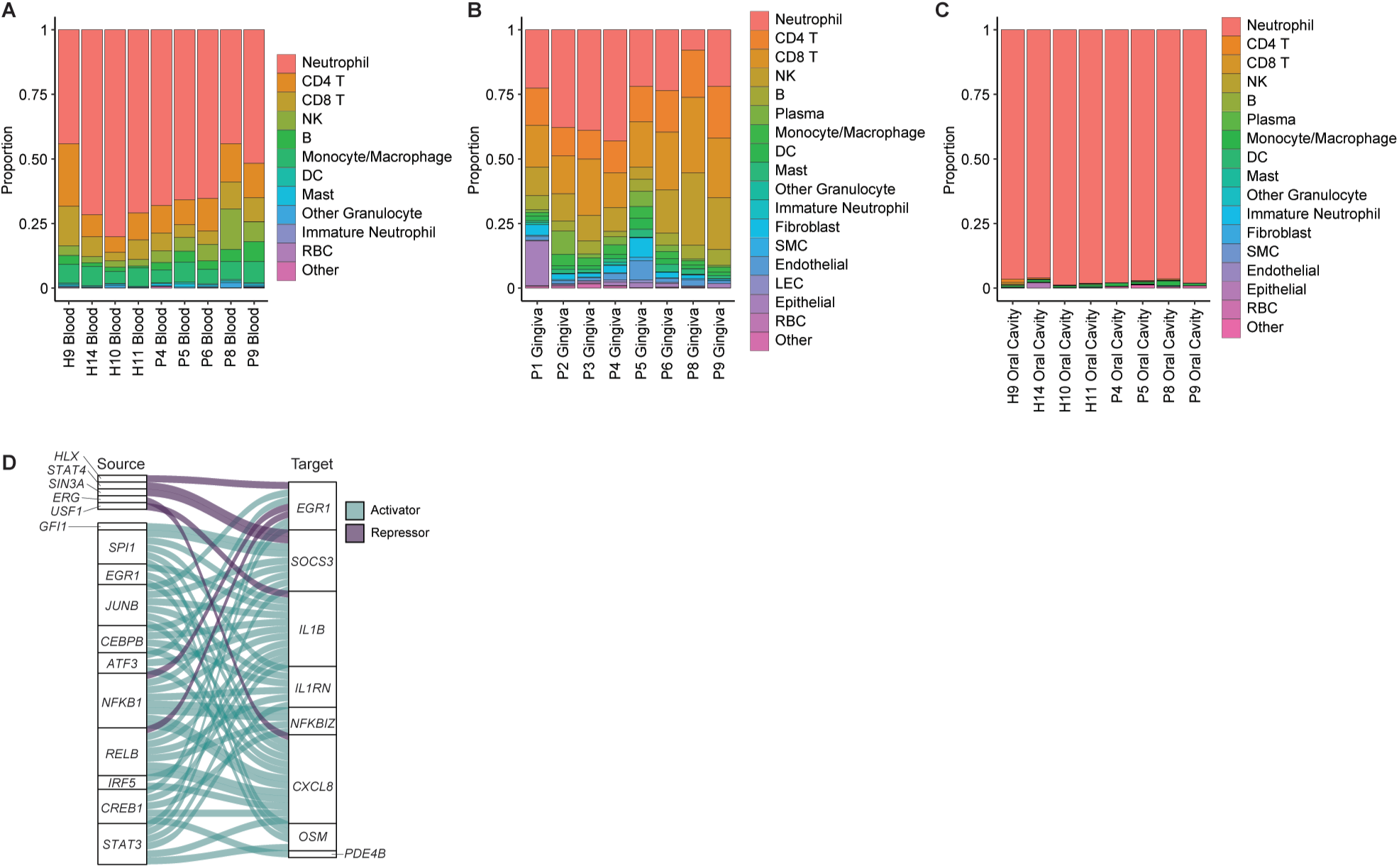
Corresponding to Figure 3. scRNA seq of immune cells across compartments. A-C) Major cell Type distribution by patient in blood (A), gingiva (B), and oral cavity (C). D) Inferred transcription factor directional activity.

**Figure S4.**
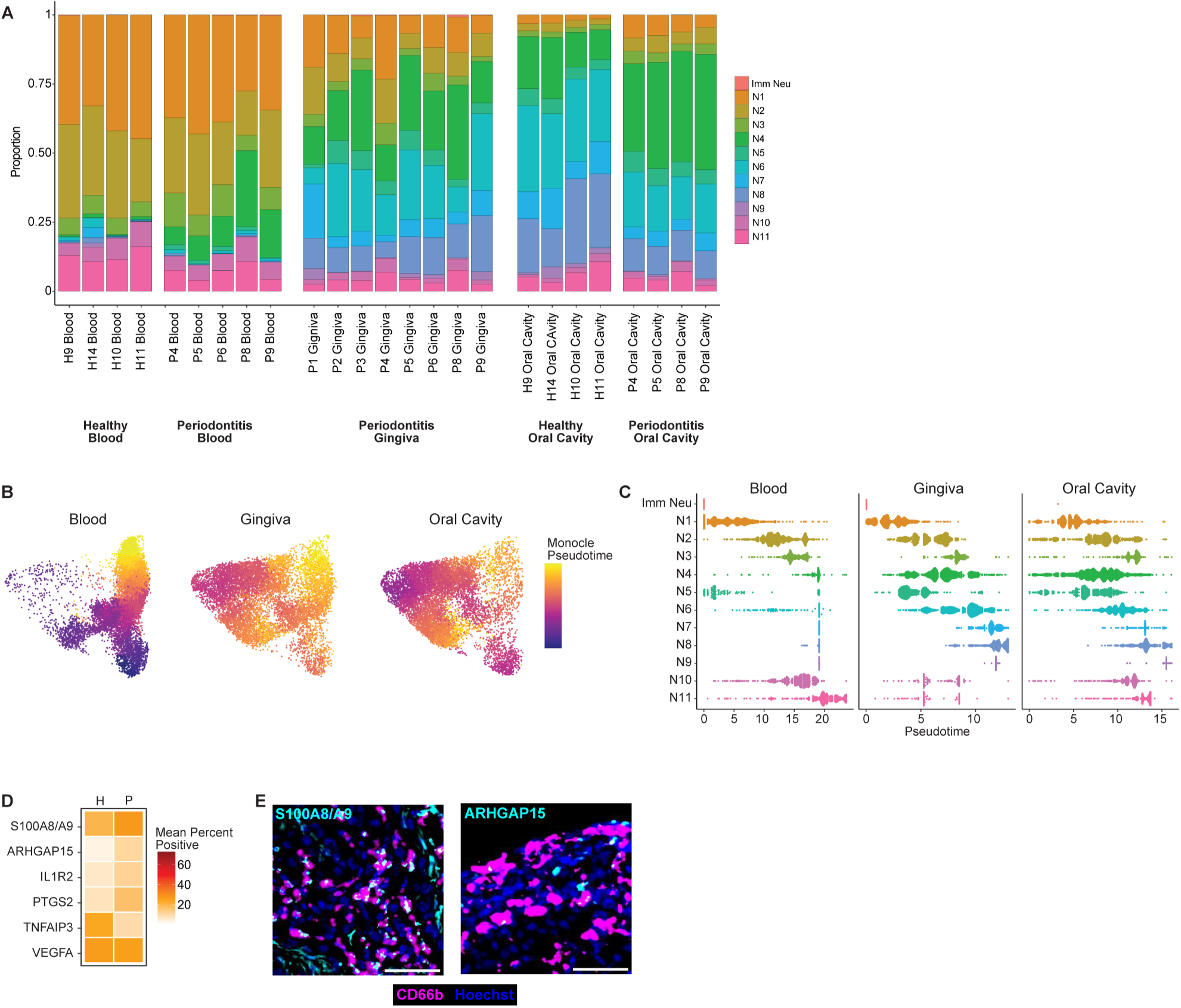
Corresponding to Figure 4. scRNA-seq of Neutrophils across compartments. A) Distribution of neutrophil clusters across individual samples. B) UMAP visualization of neutrophil pseudotime calculated using Monocle for each tissue. C) Distribution of cells by cluster across pseudotime for each tissue. D) Mean percentage of neutrophils positive for given marker in H or P gingiva. E) Representative images of neutrophils (CD66b, magenta) and S100A8/A9 (left) and ARHGAP15 (right) in cyan. Scale bars are 50 μm.

**Figure S5.**
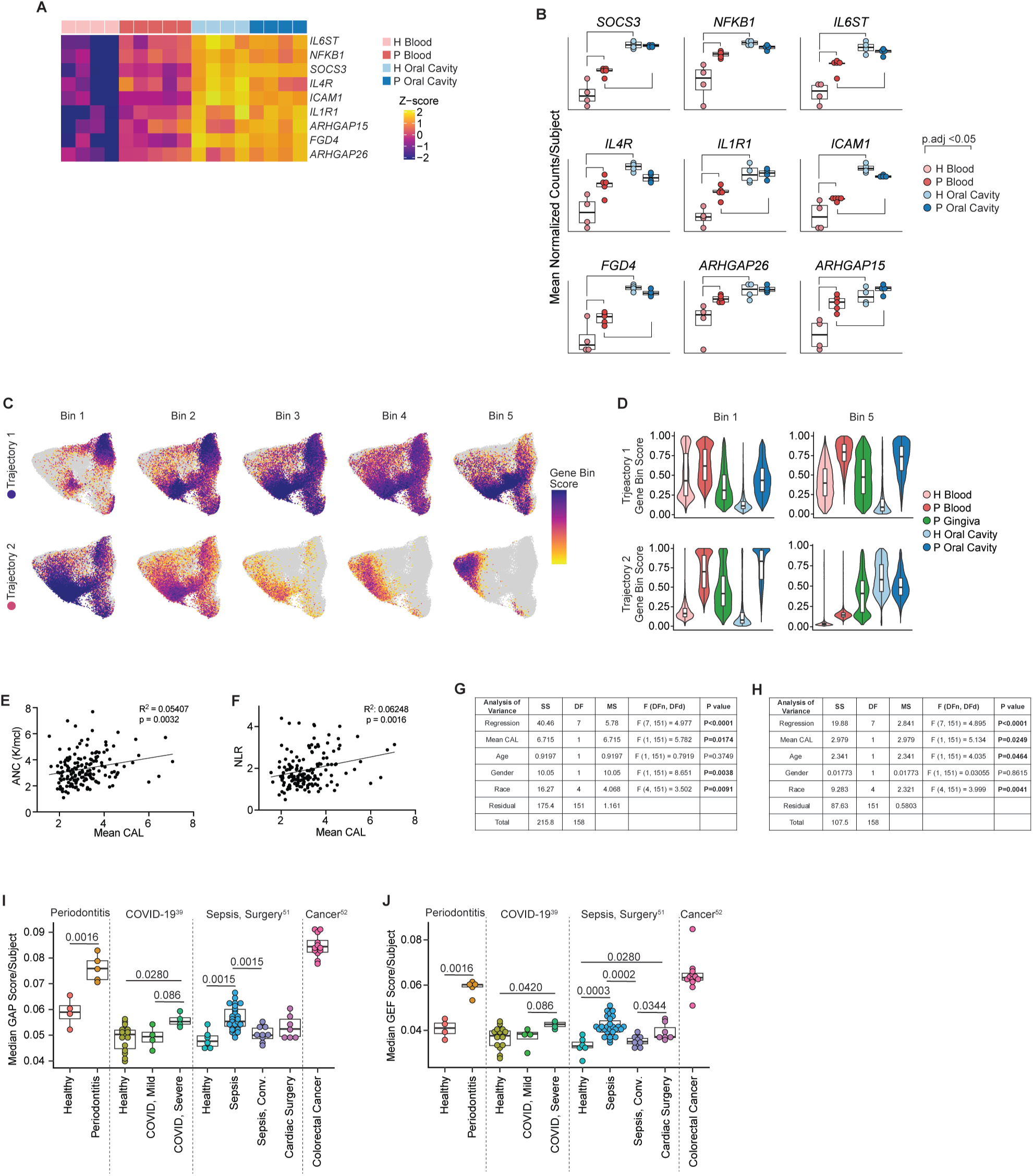
Corresponding to Figures 5 and 6. A) Scaled expression of inflammatory response and Rho-GTPase regulatory-related genes derived from P blood (Figure 6A-B) in H and P blood and oral cavity. B) Mean normalized counts by subject for genes shown in (A). Lines indicate adjusted p-values <0.05 as tested using Wilcoxon-signed rank tests with Benjamini-Hochburg adjustments for multiple comparisons. C-D) Genes in gene trajectories 1 or 2 (Figure 6F) were split into 5 even bins (Table S18). Cells were scored for expression of genes within each trajectory 1 or 2 bin and shown as a UMAP in (C) and violin plot in (D). E and F) linear regression of ANC vs. mean clinical attachment level (CAL) (E) or NLR vs. CAL (F). G and H) Results of multiple regression analysis of ANC (G) and NLR (H) and given factors. SS: sum of squares, DF: degrees of freedom, MS: mean square, F: F-test. I and J) Median GAP gene score (I) or median GEF score (J) in peripheral blood neutrophils by subject (symbol) in the current study (Periodontitis) and from studies of patients with COVID-19, sepsis or cardiac surgery, and colorectal cancer. Wilcoxon-signed rank test was used in I and J for comparisons within individual studies with Benjamini-Hochburg adjustment for multiple comparisons.

## Notes

### Competing Interest Statement

The authors have declared no competing interest.

